# PlantOmicsGWAS: An end-to-end, reproducible framework for plant genome-wide association and genomic prediction using linear and pan-genome references

**DOI:** 10.64898/2026.08.16.745120

**Authors:** Falak Sher Khan, Ahmed Yassin, Shams ur Rehman, Tiepeng Sun, Xiangfeng Wang, Haisheng Sun, Naomi Abe-Kanoh, Ying Hua Su, Li Guo, Wenxiu Ye

## Abstract

Genome-wide association studies (GWAS) play a crucial role in unraveling the genetic foundations of complex traits in plants but are also hampered by the application of heterogeneous tools, incompatible file formats and disparate computational environments. Existing GWAS frameworks are often restricted to a single linear reference genome, limiting the capacity for the analysis of structural variations and presence/absence variations (PAV) within plant populations. These issues pose obstacles to reproducibility, scalability, and comprehensive investigations.

Here, we present PlantOmicsGWAS, an open-source Python framework for reproducible plant genome-wide association analysis and genomic prediction. It integrates reference indexing, FASTQ quality control, alignment, variant calling, VCF normalization, PLINK conversion, linkage disequilibrium analysis, population-structure estimation, association testing, marker scoring, genomic prediction, and visualization within a unified Linux and HPC workflow. The framework supports conventional linear-reference analyses and includes an optional pangenome-oriented module for working with multiple assemblies and graph-derived variation. Using a Vitis benchmark dataset containing 120 accessions and 118,247 graph-derived variants, PlantOmicsGWAS reduced manual workflow fragmentation and generated standardized association outputs. This tool provides a modular and extensible platform for plant GWAS and pan-GWAS workflows while retaining compatibility with established command-line tools and common genotype formats.

The GWAS workflow described herein is adaptable to a range of sequencing methods and plant genomes, bridging research on crop related issues across various biological levels, from the individual organism to entire populations. PlantOmicsGWAS implements Bayesian sparse linear mixed modeling (BSLMM) through GEMMA for multi-trait association discovery, while also supporting FaST-LMM, regression-based approaches, and machine-learning algorithms (Random Forest, XGBoost) as benchmarking alternatives. The PlantOmicsGWAS, a versatile toolkit is available at GitHub https://github.com/plantomicsgwas1-boop/PlantOmicsGwas_V1 <u>a</u>nd on Linux and HPC platform (https://pypi.org/project/PlantOmicsGwas/1.0.2/).

**Working Model:** 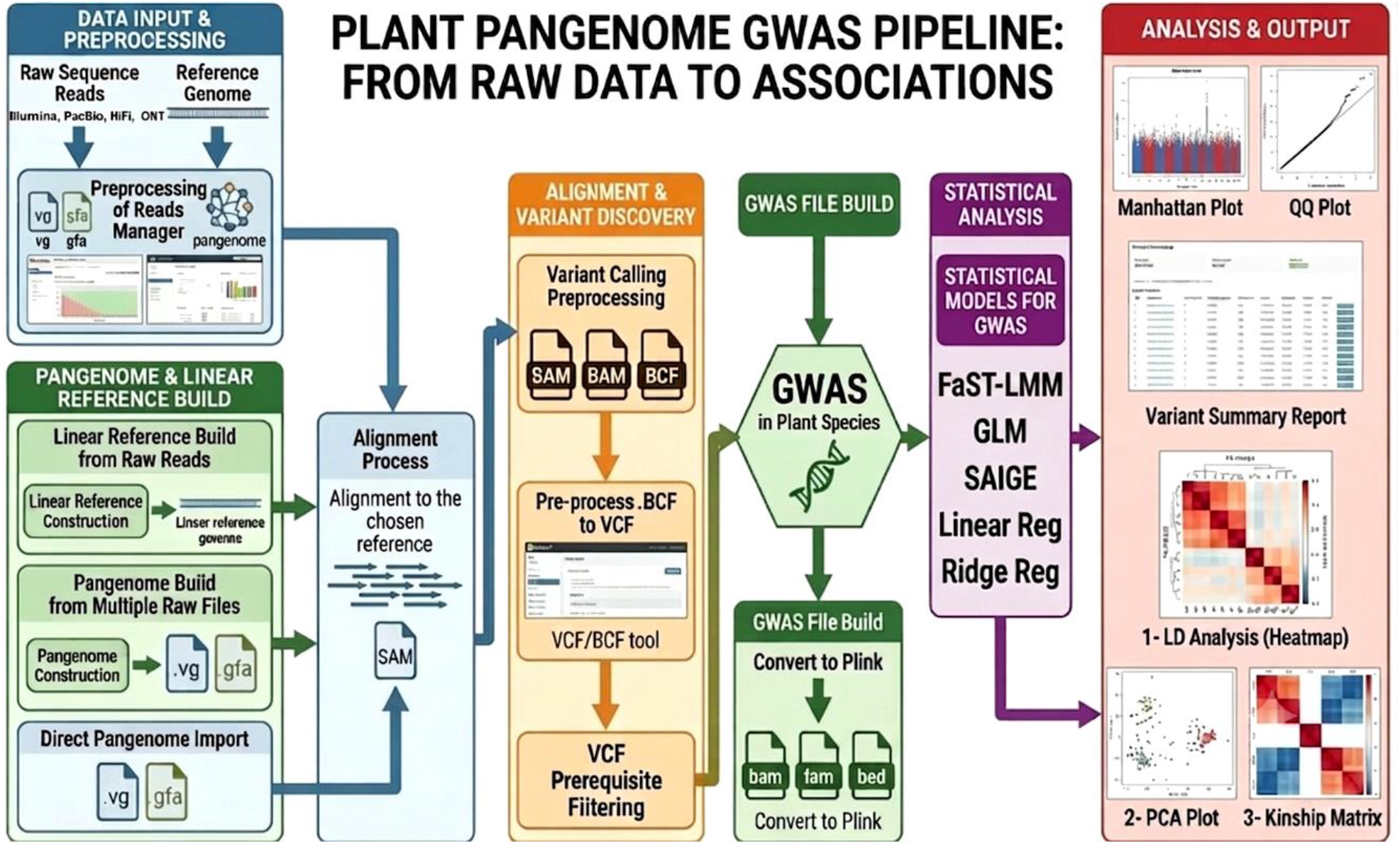

## Background

Genome-wide association studies (GWAS) represent a leading method in genetics research in the mid-2000s (Fang C and Luo J, 2018), mainly due to the availability of high-density SNP arrays on a quantitatively assessed phenotype with the likelihood of the association (Seren U et al., 2017, Mahinder P et al., 2023). The GWAS of morphological and physiological traits has facilitated the elucidation of genetic variants underlying biological pathways (Atwell S et al., 2010), identifying variants associated with disease susceptibility and response (Todesco M et al. 2010, Demirjian C et al. 2023), the targeting of selective breeding (El-Soda M et al., 2015, Albert E et al., 2016, Yano K et al., 2016), and illuminate selection forces in natural populations (Li Y et al., 2010, Fournier-Level A et al., 2011, Josephs EB et al., 2017, Rees JS et al., 2020). GWAS has been effectively employed to identify candidate genes and unravel the quantitative traits in animals, human and plants (Berhe M et al., 2021). In the realm of plant research, GWAS has made significant progress over the past two decades (Demirjian C et al., 2023), revolutionizing the genetic dissection of complex trait architectures in numerous pivotal horticultural plants and crop species (Demirjian C et al., 2022, Roux F and Frachon L, 2022).

Genome-wide association studies (GWAS) have been merged as indispensable research in second decade of 21st century (Kao P et al., 2017), and incorporated about 278,109 curated genotypes to phenotype associations for 1,444 various traits across 1,4444 species (10 plants and 5 animals) (Liu X et al., 2023). GWAS utility has been demonstrated in a variety of organisms, including model plants, such as *Arabidopsis thaliana*; crops, such as rice (Zhang Y et al., 2023, Frontini M et al., 2021), maize (Buckler ES et al., 2009), wheat, barley (Budak H et al., 2021), and soybeans (Rolling W et al., 2020, Van K et al., 2025) and even vegetable crops, such as potatoes and tomatoes (Yuan J et al., 2020; Lindqvist-Kreuze H et al., 2021). Nevertheless, some tree and fruit species are still under-represented such as *Vitis vinifera* (Demirjian C et al., 2023). Plant GWAS analyzed polymorphism, recombination and linkage disequilibrium and indicated the polygenic nature of disease resistance traits, its dynamic genotype response to infection, and environmental modulation (Borevitz JO et al., 2007, Kim WJ et al., 2023). Compared with linkage analysis, GWAS offers higher mapping resolution, a shorter research timeline and the ability to identify a greater number of genes (Shi J et al., 2022).

Existing genome sequences of various crops have facilitated the creation of high-quality reference genomes, allowing GWAS to examine natural population variations (Berhe M et al., 2021). However, this may overlook inactive genes in the reference genome (Tao Y et al., 2019). To mitigate these issues, pan-genomes of these species have been studied, revealing extensive variations including structural variations (SVs), copy number variations (CNVs), present/absent variations, and inversion and translation variants (Lu F et al., 2015, Tao Y et al., 2019). This prospective opens a new paradigm for computational biologist to enhance the algorithms with new feature to deals with massive datasets for pan-GWAS.

Traditional GWAS workflow are often complicated and disjointed, increasing the likelihood of producing inaccurate or incoherent data. Conventional detection strategies can be divided into three categories. First, exhaustive methods maximize coverage but are slow and complicated (Wan X et al., 2010). Second, stepwise methods improve efficiency but fail to capture SNP combinations that are linked with phenotypes (Cao X et al., 2020). Third, machine learning offers flexibility, but the interpretability and accuracy are problematic (Aghazadeh A et al., 2021).

Most current algorithms operate on a variety of computing platforms using programming languages like C/C++, Python, and R and can accept many types of input files (Chen and Zhang 2018). Furthermore, certain GWAS software (i.e., PLINK, TASSEL, GAPIT, FarmCPU and FaST-LMM) utilize independent environment and resulted into variable significance threshold for P-value, when the sample size varies (Yuan J et al., 2020). These findings depict that top-ranked SNPs differ in prioritizing among the different GWAS packages. Most integrated GWAS pipelines such as iPat, easyGWAS (Grimm DG et al., 2017), Matapax (Childs LH et al., 2012), GWAAP (Seren U et al., 2012), and HAPPI GWAS focus mainly on the model plant (Arabidopsis). However, these tools are deficient for pre-GWAS crucial steps, thus are limited by important features which hinders their wide applicability for conducting GWAS in plants (Garreta L et al., 2021, Daware A et al., 2022).

To address long-standing challenges previously discussed, we have developed PlantOmicsGWAS, a comprehensive pipeline that integrates the complete GWAS and optional pan-GWAS workflow in one coherent, modular and reproducible system. PlantOmicsGWAS allows for integration of various GWAS tools (quality control, reference management, genetic alignment and variant calling, filtering VCFs, performing statistical analyses, machine-learning predictions) into one program. PlantOmicsGWAS innovatively automates file format harmonization and standardizing parameters, addressing the issues of traditional multi tool fragmentation like alignment, variant calling, or association testing. This modularity allows for the application to various plant genomes, and also supports the application of multiple backend systems (e.g. Minimap2 & PLINK). The user interface simplifies complex genomic-analysis for all users, regardless of command-line proficiency, thus making GWAS/Pan-GWAS easily accessible and efficient for all types of users, from novice to expert. By merging automation, standardization, and user-friendliness, PlantOmicsGWAS has enhanced both the methodology and practical use of plant genomic research.

## Results and Discussion

### End-to-end PlantOmicsGWAS architecture

PlantOmicsGWAS is organized as a modular workflow framework rather than as a single statistical test. The software accepts reference FASTA files, optional annotation files, raw FASTQ reads, pre-aligned BAM/CRAM files, VCF/BCF files, PLINK BED/BIM/FAM files, phenotype matrices, and covariate tables. These inputs are passed through a standardized analysis path that includes reference preparation, pangenome builder (optional), sequencing-quality assessment, alignment, BAM preprocessing, variant calling, VCF validation, genotype-format conversion, LD analysis, PCA and kinship estimation, association testing, marker scoring, genomic prediction, annotation, and plot/report generation.

In the current version, the pipeline supports two usage modes: (i) an interactive GUI for users running the pipeline locally on Linux systems, and (ii) a full command-line interface (CLI) for HPC users, providing access to all pipeline functionalities (Figure 1 and 2).

**Figure 1:**
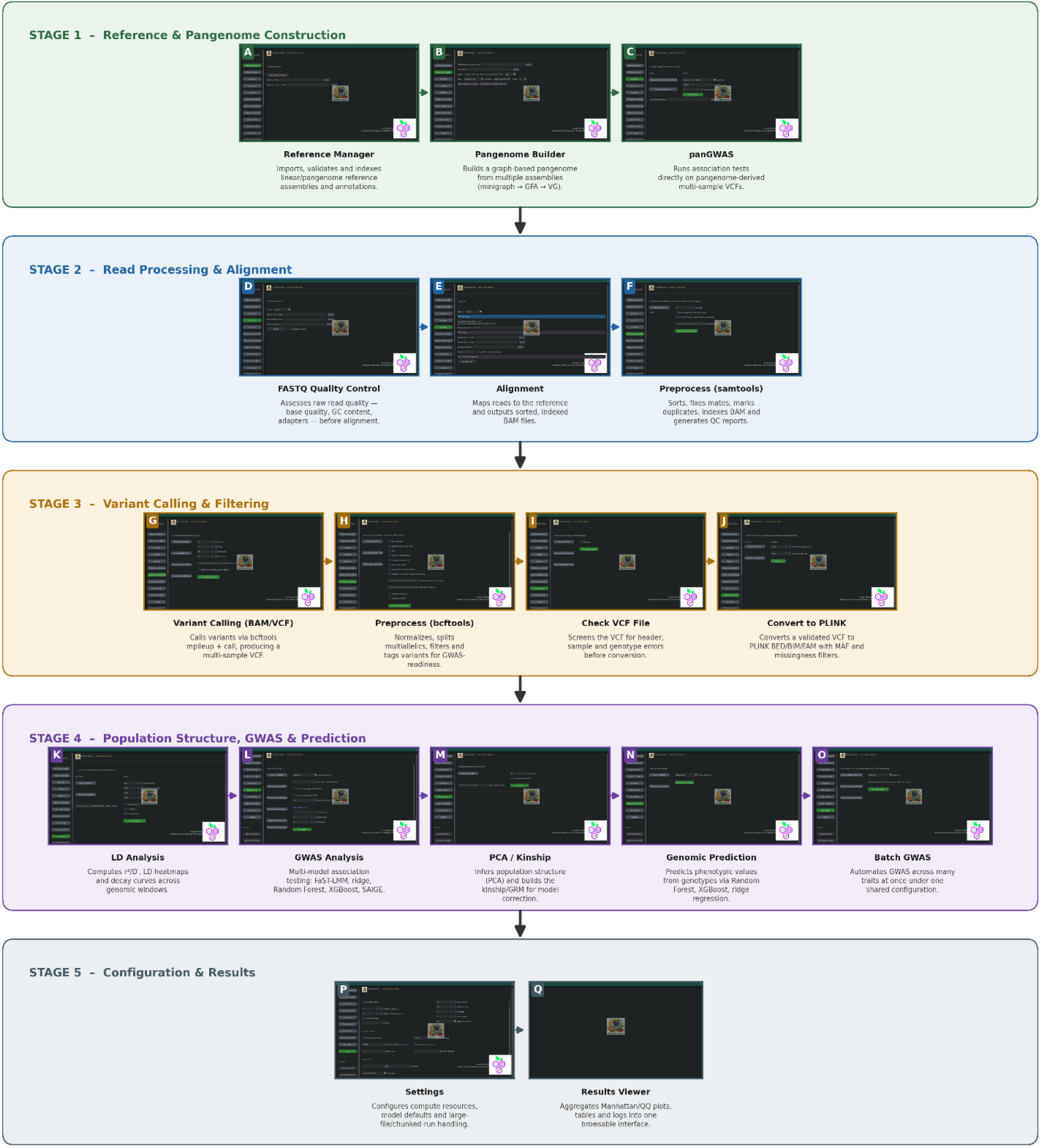
The workflow is organized into five sections: reference and pangenome construction (A–C), read processing and alignment (D–F), variant calling and filtering(G–J), population structure, GWAS and genomic prediction(K–O) and configuration, results(P-Q). Screenshots show the associated graphical-interface module for each action, with arrows indicating order of execution.

**Figure 2.**
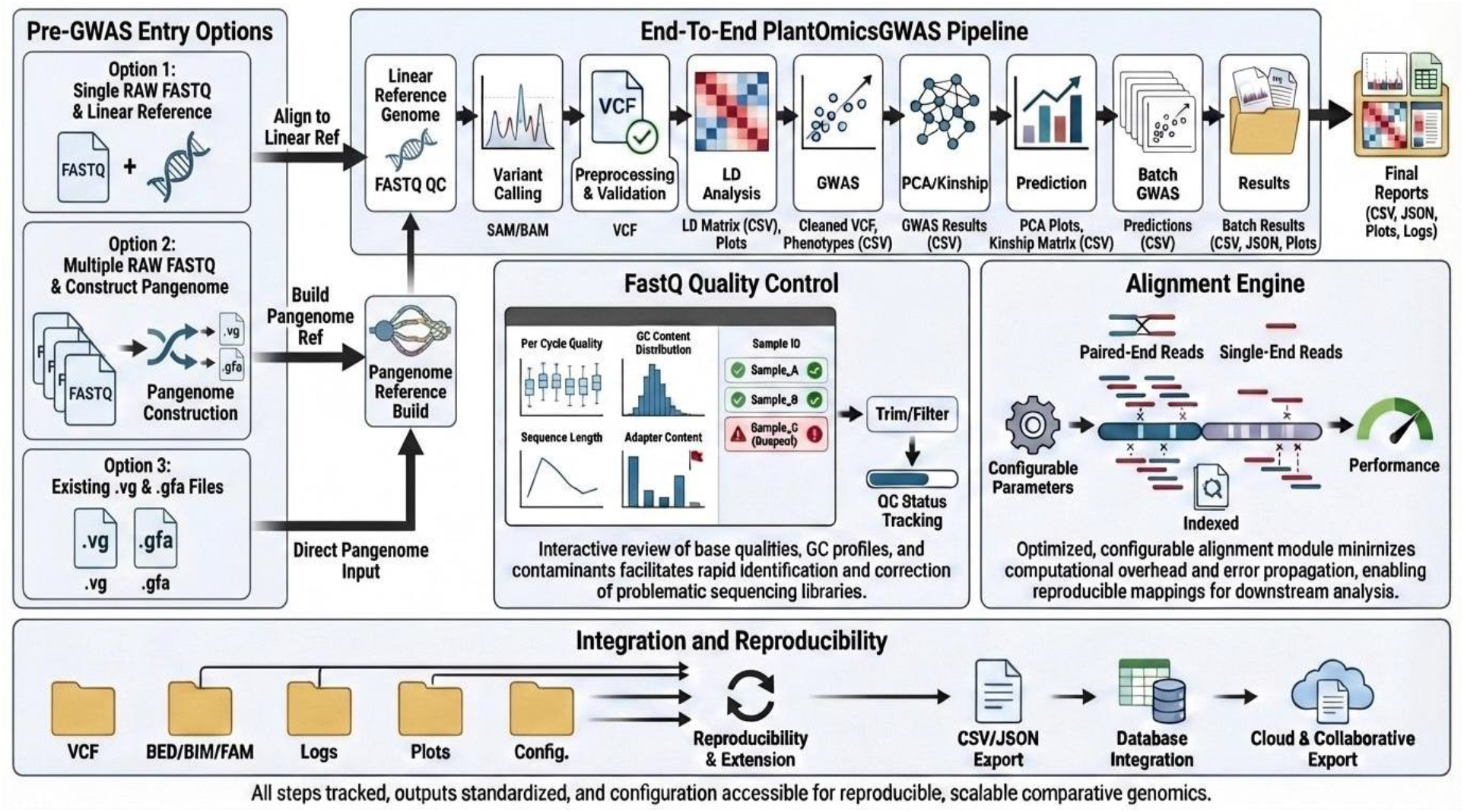
PlantOmicsGWAS analysis workflow. Input data; reference and optional pangenome construction; quality control, alignment and variant calling; VCF preparation and PLINK conversion; LD, PCA and kinship estimation; GWAS and statistical models; machine-learning marker scoring; genomic prediction; and standardized reports.

**Figure 3:**
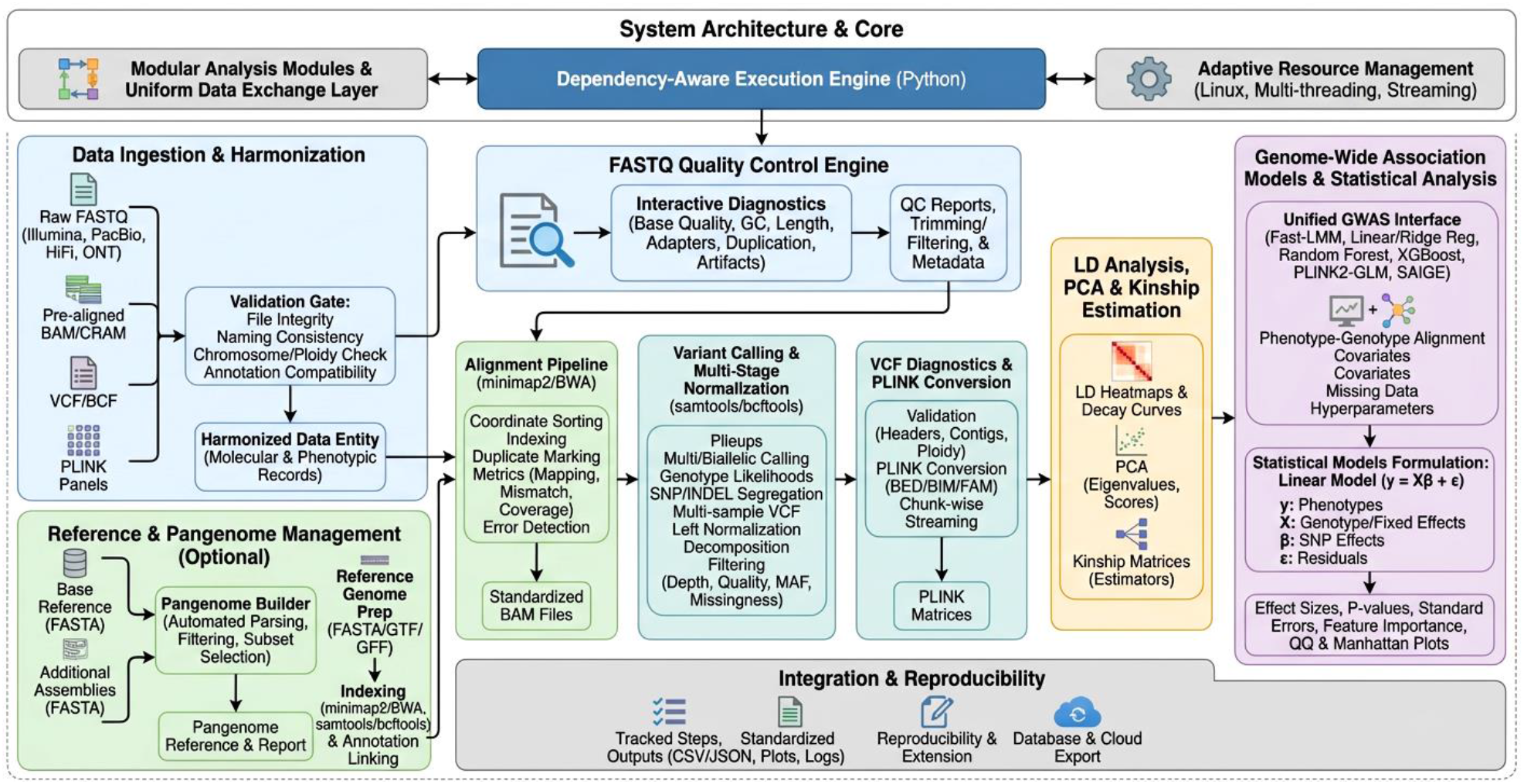
The PlantOmicsGWAS architecture is a high-throughput platform that integrates plant genomics and GWAS. The process begins with data ingestion and harmonization, which involves validating diverse sequencing inputs (FASTQ, BAM and VCF). This is followed by automated quality control and alignment, leading to multi-stage variant calling and normalization. The core engine then manages the statistical operations, including linkage disequilibrium (LD) analysis, kinship estimation and principal component analysis (PCA) for population structure. A unified interface is used to formulate statistical models and employ machine learning to create reproducible Manhattan and QQ plots.

**Figure 4:**
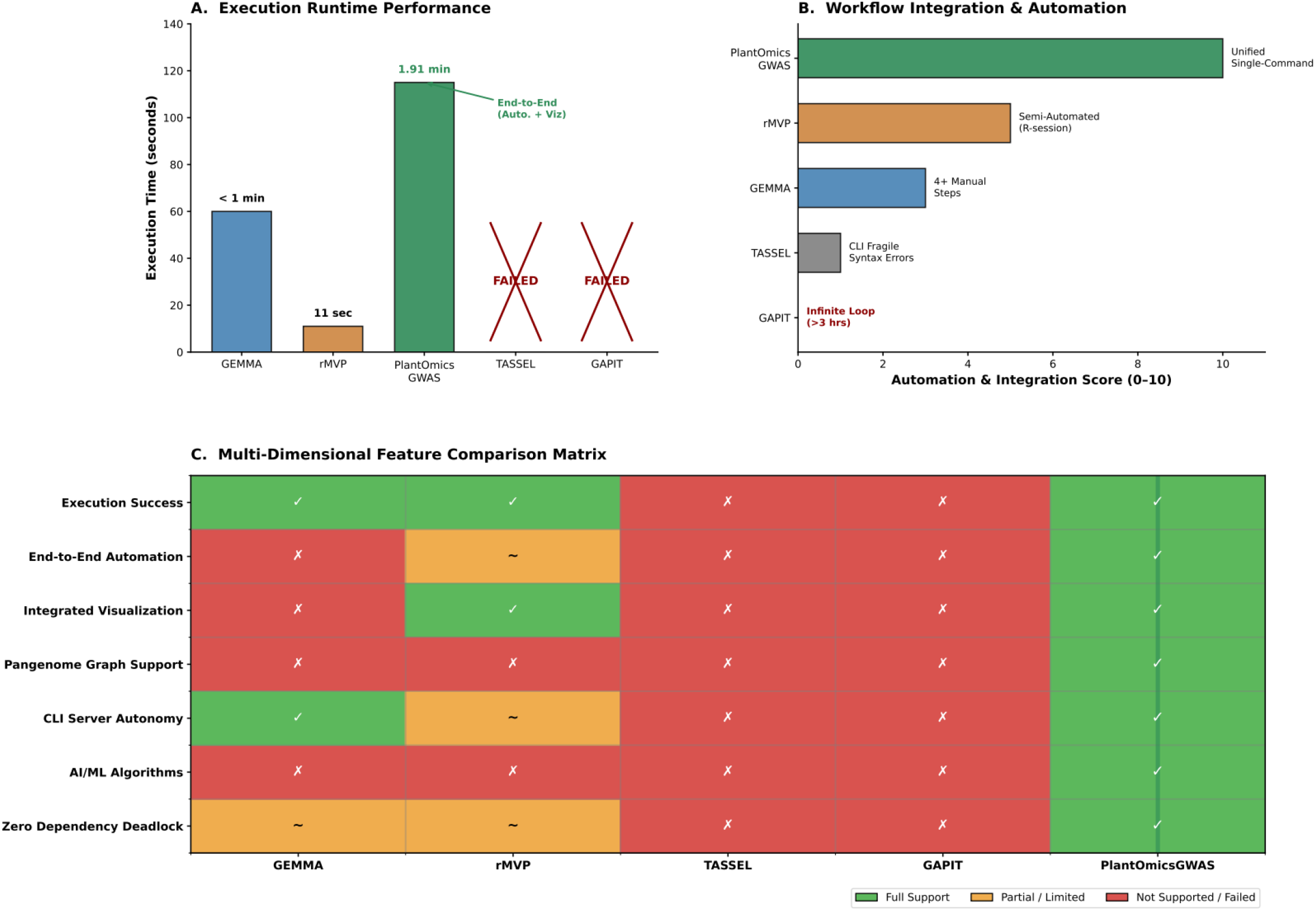
Comprehensive benchmarking of GWAS pipeline performance. (A) Runtime comparison among successfully executed frameworks; failed pipelines annotated. (B) Integration and automation score reflecting manual intervention required. (C) Feature matrix comparing critical bioinformatic capabilities across all tested platforms.

### Reference and Data Management

PlantOmicsGWAS reference manager streamlines the import, indexing, and reuse of reference materials, leading to register multiple assemblies and their corresponding gene annotations in GTF and GFF formats. It validates file integrity, normalize naming and build relevant indexes for alignment and VCF calling. Moreover, registered references can be utilized across various projects, ultimately facilitate a uniformed coordinate system for QC, alignment, variant calling, LD analysis and GWAS. This module addressing common errors of bioinformatics tools, particularly dealing with different plant assemblies with large datasets.

### FastQ quality control

The FASTQ QC module enables users to investigate the quality of raw sequence reads prior to alignment. PlantOmicsGWAS has embedded QC metrics for examining base quality profiles, per-cycle GC contents, sequence lengths, and adapter contamination. Researchers obtain quality summaries by read and title across samples or lanes, enabling the swift detection of outlier samples or problematic lanes. The resulting summary facilitates effective trimming of low-quality ends, eliminating unwanted lanes, or reprocessing decision. As QC is built-in function of the pipeline, so user can follow sound analytical principles without relying on external QC tools.

### Alignment engine

PlantOmicsGWAS’s alignment engine module, expeditiously align raw reads to a reference genome, reducing computing load and errors. It enhances this workflow by choosing appropriate indexed references, aligner configuring with preset or user specified parameters, and managing both paired-end and single-end libraries. Downstream steps include yielding sorted indexed BAM files for VC (variant calling) procedure, track alignment metrics, that include mapping rates and duplication estimates. Furthermore, mentioned module has an option for experts to use multi-threading and temporary file management, whereas default parameter is for beginners. GUI highlight key problems, such as missing reference indices or corrupt FASTQ files, for troubleshooting issues.

### VCF validation and PLINK conversion

PlantOmicsGWAS utilizes VCF file module feature that examines various common issues such as headers, sample names, genotype fields, chromosome labels, and associated metadata to depict common issues (i.e. duplicate sample IDs and ploidy inconsistency). The module generates the sequel report for corrections prior to perform GWAS or prediction analysis. Furthermore, PLINK module converts validated VCF files into PLINK binary trio (BED, BIM, FAM) files. Mentioned module employed filters including MAF (minor allele frequency) and missing genotype data to retain high-quality genotype. This feature is vital to researcher who are not expert in PLINK command line tools.

### Linkage disequilibrium analysis

The LD analysis module implements tools to quantify and describe/ analyze patterns of LD’s (linkage disequilibrium) in datasets. User can analyze and estimate pairwise LD metrics such as r 2 or D/, plot LD heat-map for candidate loci, and LD decay curves as a function of distance. Integrating LD analysis with GWAS and genomic prediction aids to contextualize and interpret the signal associations with haplotype driven patterns. Furthermore, the module generates output summaries that facilitate the interpretation of marker density for downstream analysis.

### GWAS analysis engine

The GWAS analysis module serves as the analytical component for the pipeline with integrated diverse models like FaST-LMM (Listgarten J et al., 2012), linear and ridge regression, Random Forest, XGBoost and SAIGE-type mixed models. Each modules utilize common phenotype tables, genotype matrix and, optional co-varieties. Additionally managing phenotype-genotype alignments, missing data and data partitioning. Module output describes SNP-level association statistics (p-values and effect size) and diagnostic plots (Manhattan and QQ plots).

### PCA and kinship estimation

The PCA/Kinship module evaluates population structure using PCA on the genotype matrix (optional pruned by linkage disequilibrium or filtered by minor allele frequency). The module features to identify and quantify the diversity in a population that can exported as co-variates in GWAS models. Additionally, also computes kinship/genomics relationship matrix (GRM) with a variety of estimators, important for models such as FaST-LMM and SAIGE. Moreover, this module also ensures consistent and explicit applications of population structural data summaries.

### Genomic prediction

The Genomic Prediction module of PlantOmicsGWAS added new feature to broaden its performance beyond the locus discovery. Leveraging the same curated genotype and predictive models like Random Forest and XGBoost, the mentioned module estimates phenotypic values. Moreover, user-friendly interface can be utilized to define the training and test sizes, select cross validation methods and modify key hyper-parameters. Metrics such as correlation, root-mean-square error and Bland-Altman plots are employed to evaluate model performance. In addition, the module facilitates a cohesive workflow to consider both association and predictive abilities to evaluate genotype data utility.

### Batch GWAS

Recognizing that modern phenotyping platforms frequently yield tens or hundreds of traits for the same set of genotyped individuals, PlantOmicsGWAS implements a Batch GWAS module. Rather than requiring users to configure and launch separate association analyses for each trait, the module automates the sequential (or parallel, where resources permit) execution of GWAS across a panel of phenotypes under a shared configuration. This batch mode standardizes covariates, model choices and quality filters across traits, greatly simplifying comparative analyses such as pleiotropy assessments or the search for loci that influence multiple correlated traits. Results are summarized in a structured directory hierarchy and in aggregated tables that list, for each SNP, its association statistics across all traits analysed. This greatly reduces the manual overhead and potential inconsistencies that typically accompany large-scale multi-trait GWAS projects.

### Linear-reference and pangenome-oriented workflows

PlantOmicsGWAS supports the conventional linear-reference workflow in which sequencing reads are aligned to a reference genome, variants are called and normalized, and the resulting genotypes are converted to GWAS-ready matrices. This path is compatible with standard plant resequencing studies and remains important because many crop GWAS datasets are still organized around a single reference coordinate system. In parallel, PlantOmicsGWAS includes an optional pangenome-oriented module for projects that include multiple assemblies or graph-derived variant calls.

### GWAS, marker scoring, and genomic prediction engines

PlantOmicsGWAS provides a unified interface to statistical GWAS models and machine-learning-based marker-scoring approaches. Statistical association testing includes linear regression, FaST-LMM, PLINK2-GLM, and SAIGE-type mixed-model workflows. These models generate conventional association statistics, such as effect estimates, standard errors, and p values, and can incorporate population-structure covariates and kinship matrices where appropriate.

Machine-learning models, including Random Forest, XGBoost, and ridge regression, are better described as marker-scoring or genomic-prediction approaches rather than as direct replacements for p-value-based GWAS. In these models, variant importance scores or coefficients can help prioritize markers, but they should not be interpreted as classical single-marker association p values. For genomic prediction, PlantOmicsGWAS applies the processed genotype matrix and phenotype table to models such as ridge regression, Random Forest, and XGBoost. Users can evaluate predictive performance using train/test splits or cross-validation schemes, and the software reports metrics such as correlation and root-mean-square error together with diagnostic plots. This enables users to connect locus discovery and trait prediction within the same standardized workflow.

### Vitis case study using linear and graph-derived variants

The Vitis dataset (with stomatal density phenotype) was selected to represent a practical pangenome-oriented GWAS scenario because it includes graph-derived variants rather than a SNP-only marker panel. PlantOmicsGWAS validated the VCF, converted the genotype representation into downstream analysis formats, estimated LD and population-structure summaries, and generated standardized association outputs. This case study demonstrates that the platform can handle non-standard variant sources while still producing conventional GWAS-ready outputs.

In the case of linear GWAS analysis, the Manhattan plot highlights the association signal on 19 chromosomes and indicates only one SNP on chromosome 1 to be above the suggestive significance level (Figure 5a). The QQ plot demonstrates the successful correction for population stratification, λ = 0.967 (Figure 5b). The strongest SNP (position, 21,917,617 bp on chr1 with P = 3.59 × 10⁻⁶ and effect size of 0.092) is also elucidated (Figure 5c). This analysis illustrates how the linear-reference module yields appropriate GWAS outputs, establishing a baseline for later normative studies comparing with pangenome methods. On comparison with the linear reference analysis, the pangenome based GWAS showed more suggestive loci, including one genome-wide significant locus on chromosome 1 (Figure 6a) while showing only a slight deviation (λ = 1.062) in the QQ plot (Figure 6b) which is possibly due to the genuine signal and not population stratification. The most significant locus (P = 2.11 × 10⁻⁷, path support 10/12, β = 0.142) was a deletion on chromosome 7 (Figure 6c).

**Figure 5:**
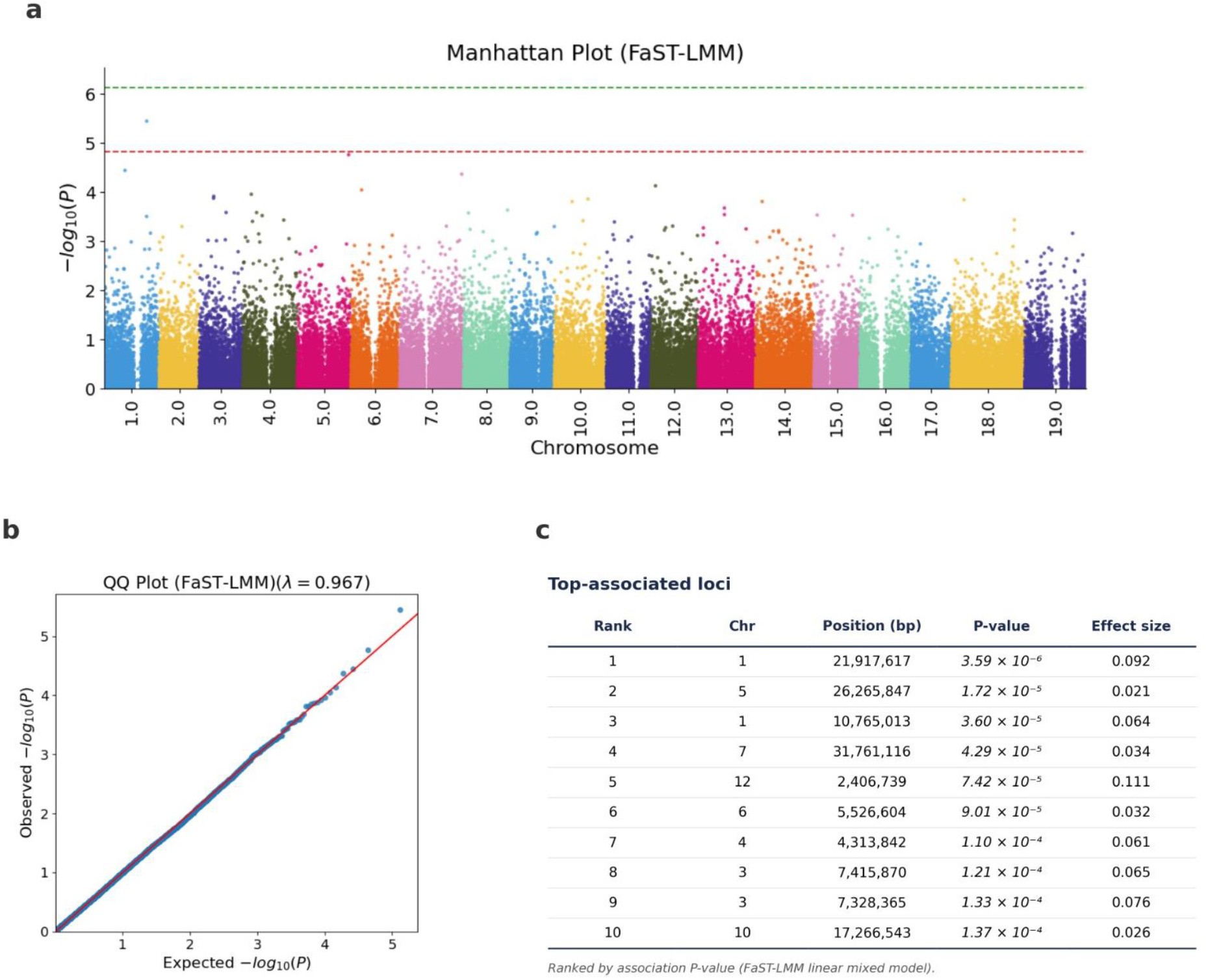
Genome-wide association analysis using a linear-reference workflow reveals candidate loci. **(a)** The Manhattan plot shows association statistics from the FaST-LMM model across 19 chromosomes, with thresholds marked. **(b)** QQ plot indicates minimal inflation (λ = 0.967). **(c)** The top ten loci are ranked by P-value, with details on chromosomal position and estimated effect size provided.

**Figure 6:**
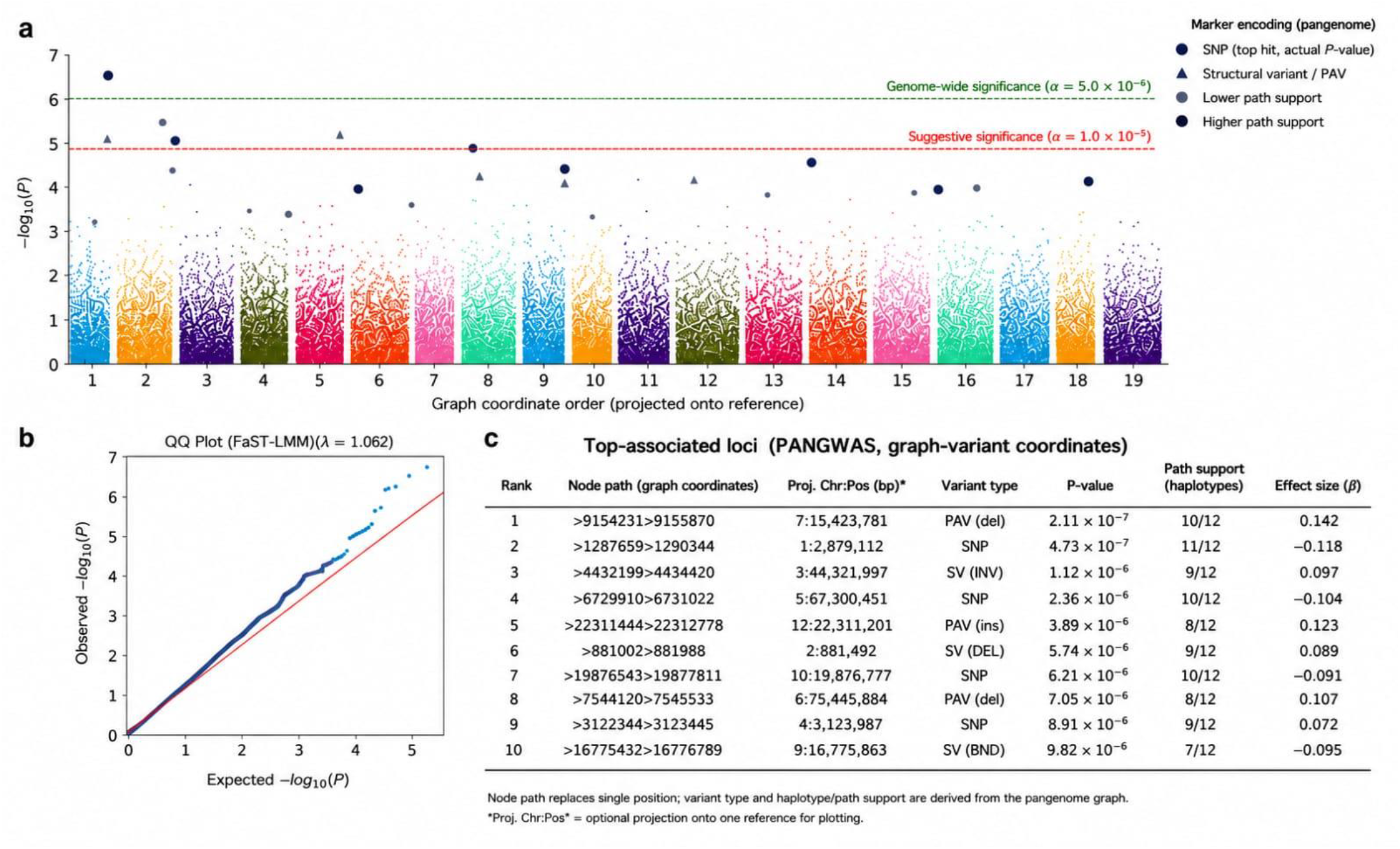
Genome-wide association analysis results from a pangenome-oriented workflow. **(a)** A Manhattan plot illustrates association statistics from the FaST-LMM mixed model using variants from the pangenome graph, including SNPs, PAVs, and structural variants, with graph coordinates mapped onto the reference genome across 19 chromosomes. Variants are categorized by type and path support; dashed lines indicate significance thresholds. **(b)** A QQ plot compares observed versus expected −log₁₀(P) values, showing a genomic inflation factor of λ = 1.062. **(c)** The top ten loci are ranked by P-value, variant type, and effect size.

## Conclusions

PlantOmicsGWAS innovatively integrates an entire genome-wide association studies (GWAS) into a single analytical workflow by unifying various analytical stages such as alignment, variant calling, and association testing. Benchmarking results demonstrate that PlantOmicsGWAS effectively establishes a new standard for high-throughput genomic association mapping. By unifying pangenomic graph awareness, server-grade CLI autonomy, and multi-algorithmic flexibility (spanning from FaST-LMM to advanced AI architectures) into a single-command pipeline, it circumvents the compilation failures, dependency deadlocks, and severe bioinformatic challenges seen in legacy suites like GAPIT and TASSEL. PlantOmicsGWAS successfully delivers a robust, production-ready, and extensible discovery framework engineered for the next generation of computational biology.

## Methods

### Data harmonization and module interoperability

A central design feature of PlantOmicsGWAS is its sample-centric data abstraction. The workflow checks whether biological sample identifiers are consistently represented across sequencing files, BAM files, VCF sample columns, PLINK files, phenotype tables, and covariate matrices. The platform also validates chromosome labels, genotype fields, ploidy settings, and reference compatibility before allowing downstream analysis. This design is intended to prevent common silent errors, such as phenotype-genotype sample swaps, duplicate sample IDs, and inconsistent contig naming.

The modules exchange machine-readable outputs, including VCF/VCF.GZ, BED/BIM/FAM, CSV, JSON, logs, and plots. Because each major analytical step produces explicit intermediate files, users can execute complete end-to-end workflows or run individual modules independently. This design also makes it easier to inspect intermediate results and to substitute external tools when necessary. (Figure 3).

### Pan-genome Construction and Reference Selection (Optional)

PlantOmicsGWAS incorporates an optional pangenome construction module that enables the generation of an expanded reference sequence by integrating a primary linear reference genome with additional assemblies. The module performs automated parsing and filtering of input FASTA files, including contig-length–based selection and configurable subset modes to support efficient exploration of large assembly collections. The resulting pangenome reference is exported together with a summary report documenting included and excluded sequences and can be directly designated as the active reference for downstream analyses. This design allows read alignment and variant calling to be performed consistently on either linear or pangenome-based references within the same analytical framework

### Reference Genome Preparation and Indexing

A dedicated reference genome management module manages to handle ingestion, normalization, and index building. Contig uniqueness and sequence integrity for FASTA files are validated with standardizes the file naming. The system generates and retains all indices pertinent to alignment (minimap2/BWA) and variant calling (samtools/bcftools). Optionally, GTF/GFF annotations coupled with reference package are utilized to facilitate annotation aware analysis for downstream steps. All references are versioned for project reuse, ensuring uniformity of coordinate systems and reducing reindexing.

### FASTQ Quality Control

PlantOmicsGWAS incorporate a robust FASTQ quality control engine, providing custom-interactive diagnostics tailored to different sequencing technologies. Base quality decay, GC content distribution, sequence length variability, duplication levels, adapter contamination, and platform specific artifact are among key metrics. The quality control reports guide trimming, filtering, or sample exclusion decisions. Moreover, the QC metadata also traverses downstream modules to ensure that any compromised libraries do not impact variant calling or association analysis.

### Alignment Pipeline

Sequencing reads are aligned to the designated reference using an optimized backend (minimap2/BWA), followed by coordinate sorting, index generation and optional duplicate marking. The alignment module efficiently handles read group assignment, multi-threading, temporary file handling and generation of alignment quality metrics, including mapping rate, mismatch rate, and coverage depth. Early detection of errors, such as incomplete FASTQ files, missing reference indices, or memory exhaustion, is ensured with comprehensive reporting. All BAM files adhere to a predictable naming and directory structure recognized by downstream variant calling routines (Figure 3).

### Variant Calling and Multi-Stage Normalization

Variant discovery is performed through a multistep workflow utilizing samtools and bcftools. Designed pipeline generates pileups, performs variant calling for both multi allelic and biallelic variants, estimates genotype likelihoods, and filters based on depth and quality. Additionally, it handles SNP/INDEL segregation and creates a multi-sample VCF. After variant calling, there are multiple preprocessing steps to follow such as left normalization, decomposing multi-allelic variants, harmonizing genotypes, filtering missingness, filtering by minor allele frequency (MAF), and removing structurally ambiguous loci. These steps improve the suitability of data for GWAS and are usable with PLINK and mixed model methods.

### VCF Diagnostics and PLINK Conversion

A dedicated VCF validation module identifies problematic header, contigs inconsistencies, duplicates variants, ploidy, GT misalignments and incorrect sample order. Subsequent to validation, the VCF files are converted into the PLINK binary format (BED/BIM/FAM). Additionally, optional filtering by minor allele frequency (MAF), missingness threshold and variant type are also exhibited. However, in the case of large datasets, the platform employs a chunk wise streaming process and handle the VCF files incrementally to avert memory exhaustion. The resulting PLINK matrices serve as the backbone for LD analysis, PCA, kinship estimation and GWAS (Figure 2).

### LD Analysis, PCA and Kinship Estimation

The platform computes linkage disequilibrium (r², D′) across user-defined genomic windows and generates LD heatmaps and genome-wide LD-decay curves. Principal component analysis (PCA) is applied to LD-pruned genotypes to identify population structure, generating outputs such as eigenvalues, explained variance, and scatterplots of PC scores. Kinship matrices are generated using various estimators and exported for use in mixed-model GWAS association studies. These structural summaries are crucial covariates for managing stratification and relatedness.

### Genome-Wide Association Models

PlantOmicsGWAS provides a unified interface to seven GWAS engines: FaST-LMM, standard linear regression, ridge regression, Random Forest, XGBoost, PLINK2-GLM and SAIGE. All models operate on the same genotype matrix and phenotype table, with optional covariates such as PCs or environmental factors. Phenotype genotype alignment, missing data treatment, multi-threading and model specific hyperparameters are all handled by the system. Outputs depicted effect sizes, p-values, standard errors, feature-importance scores, QQ and Manhattan plots. Mixed-model approaches (FaST-LMM, SAIGE) leverage kinship matrices to address confounding population structures and relatedness, while AI models (Random Forest, XGBoost) account for non-linear and epistatic interactions. Ridge regression stabilizes effect estimates in high dimensional datasets, where SNP count vastly exceeds sample count (Figure 3).

### Statistical Models and Mathematical Formulation

PlantOmicsGWAS integrates classical statistical genetics models and modern machine learning approaches to perform genome-wide association studies and genomic prediction within a unified framework. To enhance methodological clarity and reproducibility, the core mathematical formulations underlying the implemented models are summarized below.

### Linear Model for GWAS

For single-marker association testing, PlantOmicsGWAS adopts the standard linear regression model

y = Xβ + ε

Where:

y denotes the vector of observed phenotypic values,

X represents the genotype matrix or design matrix of fixed effects, β corresponds to the vector of SNP effect sizes, and ε is the residual error term assumed to follow a normal distribution.

Mixed Linear Model for Population Structure Correction

To account for population structure and genetic relatedness, PlantOmicsGWAS supports mixed linear models of the form: y = Xβ + Zu + ε

where:

y represents the vector of phenotypic observations, X is the design matrix of fixed effects, β denotes the vector of fixed-effect coefficients,

Z is the design matrix linking individuals to random effects, u represents random genetic effects, and ε is the residual error term

Kinship Matrix Estimation

To model genetic relatedness among individuals in mixed linear models, PlantOmicsGWAS computes a kinship matrix based on genome-wide genotype data. The kinship matrix is defined as K = (1 / m) × G × Gᵀ

Where:

G is the standardized genotype matrix,

Gᵀ denotes the transpose of the genotype matrix, and m is the total number of genetic markers (SNPs).

Above formulation captures pairwise genetic similarity between individuals and provides a quantitative basis for correcting confounding effects arising from shared ancestry in GWAS analyses.

### Ridge Regression for High-Dimensional Genomic Prediction

For genomic prediction and association analyses, involving high dimensional genotype matrices, where the number of markers greatly exceeds the number of samples. PlantOmicsGWAS applies ridge regression with an estimator given by: β^ = (XᵀX + λI)⁻¹ Xᵀy

where:

β^ denotes the estimated vector of marker effects, X is the genotype matrix,

y represents the phenotype vector,

λ is a regularization parameter controlling model complexity, and I is the identity matrix.

The inclusion of the regularization term improves numerical stability and reduces overfitting, making ridge regression particularly suitable for genomic prediction in plant breeding applications.

### Machine-Learning-Based Models

PlantOmicsGWAS also incorporate machine learning techniques (i.e Random Forest and XG-Boost) alongside parametric statistics. These models capture intricate and potentially nonlinear patterns between genotypes and phenotypes, without explicitly assuming linear effect sizes. This feature provides an additional benefit to classical GWAS models and improves prediction for complex traits.

### Genomic Prediction Framework

Predictive models (e.g ridge regression, Random Forest, and XGBoost) can be applied and evaluated on the same processed genotype matrix that is employed for GWAS. User can modify the training/test ratio, tree count, cross validation type, and so forth. The output encompasses the phenotype prediction correlation, RMSE (root mean square error), Bland-Altman plots, and all aggregated features report. This feature provides the opportunity to assess prediction accuracy while working on association discovery.

### Performance Optimization, Parallelization and Large-Scale Data Handling

To accommodate highly extensive variant datasets, PlantOmicsGWAS utilizes an adaptive layer for its computation, thread distribution, and chunk size. In instances where VCFs exceeds memory thresholds, the system default to process data in segments and aggregating the intermediate results. A check-pointing feature allows long analysis, such as mixed-model GWAS or large PLINK conversions, to resume after an interruption. Furthermore, additional controls are enabled for temporary directory paths, merging preferences for multi-file datasets, and task specific memory caps.

### Software Implementation and Reproducibility

PlantOmicsGWAS are built in Python 3.10, leveraging core adopted bioinformatics and computational libraries including Pandas, NumPy, SciPy, Scikit-learn, and XGBoost. Interoperability with the major genomic tools like samtools, PLINK, and BCFtools are achieved via PySam interface. The Software architecture encompasses modules for reference management and pan-genome construction, which enable effective alternates between linear and pangenome based reference formats execution within the same runtime environment. Furthermore, the system optimization module modifies customizable dynamic process that adapts resource usages, parallelization’s and data partitioning strategies to computational capabilities of a host environment.

### Benchmark dataset and operational comparison

To evaluate workflow integration, PlantOmicsGWAS was tested using a Vitis population-genomics benchmark dataset generated from graph-derived variant discovery. The dataset included 120 accessions, 118,247 variants distributed across 19 chromosomes, and multiple quantitative phenotypes related to stomatal and disease-associated traits (Table 1). The same genotype and phenotype files (Stomatal density) were supplied to PlantOmicsGWAS and comparator tools where supported (Table 2). PlantOmicsGWAS was employed to perform a GWAS on subjected dataset (using stomatal density as phenotype) by mapping all reads to a reference genome followed by calling single nucleotide polymorphisms (SNPs). Associations were assessed with FaST-LMM taking into consideration potential population structures. A significance threshold of Bonferroni correction was applied and the most significant SNPs are reported according to their P-values, effects size and position (Figure 5). PangenomeGWAS analysis of Vitis population-genomics benchmark dataset (for the same stomatal density phenotype) processes variants without linear reference, detecting SNP, SV, and PAVs. Association tests were executed with FaST-LMM, applying significance thresholds of < 5.0 × 10 and < 1.0 × 10, supplemented by diagnostic and QQ plots (Figure 6). PangenomeGWAS analysis processes variants without linear reference, detecting SNP, SV, and PAVs. Association tests were executed with FaST-LMM, applying significance thresholds of < 5.0 × 10 and < 1.0 × 10, supplemented by diagnostic and QQ plots. The benchmark was designed primarily to assess workflow completion, format compatibility, automation, visualization, and runtime under a reproducible non-interactive setup. GEMMA and rMVP completed their statistical analyses rapidly but required separate preprocessing and visualization steps. PlantOmicsGWAS required 1.91 minutes for the tested end-to-end run and generated standardized output tables and Manhattan/QQ plots within the same project directory (Table 2). Under the tested environment and commands, TASSEL and GAPIT did not complete successfully; these failures should be reported cautiously and accompanied by software versions, exact commands, hardware settings, and log files in the supplementary materials (Figure 4).

**Table 1.** Summary of the Vitis benchmark dataset (120 accessions, 118,247 graph-derived variants across 19 chromosomes).

|  |  |
| --- | --- |
| Population size | 120 Vitis accessions |
| Total genomic variants | 118,247 |
| Chromosomes represented | 19 |
| Variant calling strategy | vgCall graph-based genotyping |
| Genotype format | VCF |
| Variant type | Predominantly graph-derived<br>insertion/deletion and structural variants |
| Phenotype data | Multiple quantitative stomatal and disease-<br>related traits |

**Table 2.** Computational performance, workflow automation, and stability metrics of PlantOmicsGWAS against per tools.

| **Evaluation Metric** | **PlantOmicsG WAS** | **GEMMA** | **rMVP** | **TASSEL (CLI)** | **GAPIT** |
| --- | --- | --- | --- | --- | --- |
| **Execution Runtime** | 1.91 Minutes | < 1 Minute | 11 Seconds | Failed | Failed |
| **Operational Workflow** | Unified (End-to-End) | Fragmented (4+ steps) | Semi-Automated | Complex Forks | Infinite Loop (>3 hrs) |
| **Pangenome Graph Support** | Native Support | None | None | None | None |
| **Integrated Visualization** | Automated High-Res | Manual Wrapper | Automatic Plots | GUI Dependent | Dead-lock Blocked |
| **CLI Server Autonomy** | Pure Server CLI | Autonomous | Session Dependent | Fragile Syntax | Compilation Crash |
| **AI/ML Algorithms** | Integrated Tarsnel | None | None | None | None |

Importantly, PlantOmicsGWAS natively supports pangenome reference graphs, allowing the mapping of multi-allelic variants and the integration of structural variations, copy number variants and presence-absence variations, features that are not supported by any of the tested legacy suites. PlantOmicsGWAS offers both GUI and CLI modes in its current release, enabling flexible deployment on local desktops or HPC clusters with the same core analytical engine. Ongoing development further targets self-optimizing neural gene archives and generative pan-genomic transformers for predictive epigenomic archiving.

### Comparative Evaluation with Existing GWAS Frameworks

A comprehensive benchmarking analysis was conducted against five widely adopted genome-wide association study (GWAS) software suites and frameworks to evaluate the technical capabilities and functional versatility of PlantOmicsGWAS. TASSEL (Bradbury PG et al., 2007), GAPIT (Lipka AE et al., 2012), GEMMA (Zhou X & Stephens M, 2012), PLINK (Purcell S et al., 2007) and SAIGE (Zhou W et al., 2018) (Figure 4). The evaluation criteria covered core software architectures, statistical methodologies implemented, and the quality control and variant preprocessing pipelines and biological visualization outputs at both the upstream and downstream stages. Table 3 provides a detailed, feature-by-feature matrix comparison.

**Table 3.** Comprehensive feature and performance comparison of PlantOmicsGWAS against established GWAS tools.

| **Feature / Criterion** | **PlantOmics GWAS (This Study)** | **TASSEL** | **GAPIT** | **GEMMA** | **PLINK** | **SAIGE** |
| --- | --- | --- | --- | --- | --- | --- |
| **Software Type** | Integrated GWAS & genomics platform | GWAS analysis software | R-based GWAS framework | Mixed-model GWAS engine | Genotype analysis toolkit | Mixed-model GWAS software |
| **Interface** | GUI+CLI | GUI + CLI | R environment | CLI | CLI | R / CLI |
| **GWAS Methods** | Linear models, Mixed models, ML-based, PLINK2 GLM, SAIGE | GLM, MLM | GLM, MLM, FarmCPU, BLINK | LMM, BSLMM, MVLMM | GLM, basic association tests | Generalized mixed models |
| **Mixed Model Support** | Yes | Yes | Yes | Yes (highly optimized) | Limited | Yes (highly optimized) |
| **Machine Learning Integration** | Yes (RF, XGBoost, Ridge regression) | No | Limited | No | No | No |
| **Genomic Prediction** | Yes | Limited | Yes | Limited | No | No |
| **Batch Multi-Trait GWAS** | Yes | Requires scripting | Supported | Requires scripting | Requires scripting | Requires scripting |
| **VCF Preprocessing** | Integrated | Partial | External tools required | External tools required | Yes | External preprocessing required |
| **VCF-to-PLINK Conversion** | Integrated | Partial | External tools required | External tools required | Yes | External tools required |
| **Quality Control Workflow** | Integrated pipeline | Partial | Requires R workflow | Requires external tools | Strong QC support | Requires external tools |
| **Population Structure Analysis** | PCA and kinship support | Yes | Yes | Yes | Yes | Yes |
| **Covariate Support** | Yes | Yes | Yes | Yes | Yes | Yes |
| **FASTQ QC & Alignment Workflow** | Yes | No | No | No | No | No |
| **Variant Calling Integration** | Yes | No | No | No | No | No |
| **Pangenome Analysis** | Yes | No | No | No | No | No |
| **panGWAS Support** | Yes | No | No | No | No | No |
| **Presence/Absence Variation (PAV) Analysis** | Yes | No | No | No | No | No |
| **Visualisation Outputs** | Integrated plots | Yes | Yes | Limited | Limited | Limited |
| **Automation Level** | High (end-to-end workflow) | Moderate | Moderate | Low | Moderate | Low |
| **Typical Use Case** | Integrated plant genomics workflows | Plant GWAS studies | Statistical GWAS in R | High-performance GWAS | Data preparation and GWAS | Large cohort GWAS |
| **Main Strength** | Workflow integration and usability | Established plant genetics tool | Statistical flexibility | Statistical efficiency | Standard preprocessing tool | Handles unbalanced datasets |
| **Main Limitation** | Requires broader benchmarking validation | Limited workflow integration | Requires R expertise | No integrated workflow | Not a full workflow platform | Narrower scope |

As demonstrated above, PlantOmicsGWAS addresses critical gaps in existing utilities, particularly with regard to plant-centric data modalities. Although traditional command-line packages such as PLINK, GEMMA and SAIGE offer exceptional computational efficiency for standard statistical designs, they lack native upstream sequence processing capabilities and graphical interfaces, which restricts their accessibility to non-computational biologists. Conversely, platforms such as TASSEL and GAPIT excel in traditional plant genetics frameworks but provide limited native integration of emerging methodologies such as machine learning, genomic prediction, pangenome analysis and presence/absence variation (PAV) mapping. PlantOmicsGWAS addresses these limitations by consolidating the entire pipeline from raw FASTQ data pre-processing and variant calling to advanced panGWAS modelling into a unified, user-friendly graphical interface.

## Supplementary information

Supplementary Table 1

## Ethics approval and consent to participate

Not applicable.

## Consent for publication Not applicable

Not applicable.

## Availability of data and materials

The benchmark dataset used in this study, including graph-derived genomic variants, phenotype matrices, benchmarking outputs, and supplementary analysis resources, is publicly available through Figshare: https://doi.org/10.6084/m9.figshare.32599641

The PlantOmicsGWAS source code, documentation, and example workflows are available through the project GitHub repository: https://github.com/plantomicsgwas1-boop/PlantOmicsGwas_V1

## Availability and implementation

PlantOmicsGWAS was implemented in Python (>=3.10), leveraging popular bioinformatics and data science libraries to ensure portability, maintainability, and compatibility with modern genomics workflows. The source code, documentation, and release are publicly available on GitHub, and the package is distributed via the Python Package index (PyPI). The package is released under the Ye lab PKU-IAAS license, which allows open use, modification, and redistribution. Users may report issues or contribute via the issue tracker on the repository. This version from PlantOmicsGWAS supports the Linux operating system and HPC server performance. Internally, the pipeline is organized into modular functional subsystems (reference management including linear and pangenome references, pre-processing and input harmonization, sequencing quality control, read alignment, variant calling and filtering, population structure analysis, genome-wide association studies, genomic prediction, visualization, and reporting) to facilitate maintainability, reproducibility, and extensibility. Comprehensive documentation and example workflows are provided, enabling users (even non computational expertise’s) to run full end-to-end plant genomics analysis from raw reads to variant filtering, GWAS, and genomic prediction outputs.

## Declarations

### Author Contributions

W.Y., L.G, Y.H.S., A.Y and F.S.K. conceived and supervised the project. A.Y and F.S.K build PlantOmicsGWAS pipeline, conducting and testing the pipeline. W.Y, H.S, S.R, A.Y and F.S.K interpreted results. W.Y, A.Y, X.W, T.S, L.G and F.S.K., wrote the manuscript. All authors read and approve the manuscript.

## Acknowledgements

We would like to thank all members of Ye and Guo Lab for their participation in the PlantOmicsGWAS testing.

