## Supplemental Table 1 for "PlantOmicsGWAS: An end-to-end, reproducible framework for plant genome-wide association and genomic prediction using linear and pan-genome references"

**PlantOmicsGWAS (PlantOmicsGwas 1.0.2) Run Performance Report**

Execution environment metadata, resource metrics, and pipeline benchmark logs.

**Supplementary Table 1: PlantOmicsGWAS (PlantOmicsGwas 1.2.10) Execution Environment and Performance Metrics**

| Category | Parameter / Step | Details / Value |
|---|---|---|
| Run Metadata | Run ID | 2026-07-13_14-32-01 |
| Run Metadata | Start Time | 2026-07-13 14:32:01 UTC |
| Run Metadata | End Time | 2026-07-13 15:47:22 UTC |
| Run Metadata | Total Wall Time | 01:15:21 |
| Run Metadata | Exact Command | plantomicsgwas run --config config.yaml --module gwas |
| Software Environment | Operating System | Ubuntu 22.04.3 LTS |
| Software Environment | Job Scheduler | SLURM 22.05.9 |
| Software Environment | Python Version | 3.11.6 |
| Software Environment | PlantOmicsGWAS | v1.2.10 |
| Software Environment | NumPy | 1.26.4 |
| Software Environment | Pandas | 2.2.1 |
| Software Environment | scikit-learn | 1.4.0 |
| Software Environment | statsmodels | 0.14.1 |
| Software Environment | GEMMA | 0.98.5 |
| Hardware | Node Name | compute-node-07 |
| Hardware | CPU Model | Intel Xeon Gold 6248R |
| Hardware | Allocated CPU Cores | 24 cores |

| | | |
|---|---|---|
| Hardware | Allocated RAM | 64 GB |
| Hardware | GPU | None |
| Resource Usage (Peak) | CPU Usage | 92% |
| Resource Usage (Peak) | Memory Usage | 41.2 GB / 64 GB |
| Resource Usage (Peak) | Disk I/O | 3.4 GB read / 1.1 GB written |
| Pipeline Steps & Timing | Step 1: Data Loading & QC | Start: 14:32:01<br>End: 14:34:15<br>Duration: 00:02:14 |
| Pipeline Steps & Timing | Step 2: Kinship Matrix Computation | Start: 14:34:15<br>End: 14:43:02<br>Duration: 00:08:47 |
| Pipeline Steps & Timing | Step 3: GEMMA-BSLMM Association | Start: 14:43:02<br>End: 15:41:35<br>Duration: 00:58:33 |
| Pipeline Steps & Timing | Step 4: Output Aggregation | Start: 15:41:35<br>End: 15:47:22<br>Duration: 00:05:47 |
